# Robust genome-based delineation of bacterial genera

**DOI:** 10.1101/2025.03.17.643616

**Authors:** Charlie Pauvert, Thomas C.A. Hitch, Thomas Clavel

## Abstract

**Background:** Genomic analysis has become essential in bacterial taxonomy, enabling fast classification of bacteria. However, quantifiable measures to make informed decision on the taxonomic placement of bacterial taxa above the level of species are rare, which hampers the stability of knowledge in databases and articles. In this work, we focused on bacterial classification at the genus level and revisited the concept of Percentage Of Conserved Proteins (POCP). Whilst POCP is broadly used as an overall genome relatedness index, the underlying tool and method of calculation differ, resulting in disparate implementations and misnomers unfit for the increasing wealth of public genomes available.

**Results:** We evaluated 10 protein alignments methods and found evidence for a scalable yet robust alternative to BLASTP for POCP based on 2,358,466 pairwise comparisons of 4,767 genomes across 35 families. However, we showed that certain combinations of tools and parameters are not suited for POCP calculations owing to drastic over-or underestimation. Therefore, we bring forward a clearer definition of POCP using only unique matches, termed POCPu, that showed better genus delineation than with POCP. We suggested tentative family-specific thresholds when the standard of 50% was not resolutive enough.

**Conclusions:** We propose that faster bacterial genus delineation should be calculated with DIAMOND using very-sensitive settings and only unique matches (POCPu). We advise microbiologists to carefully assess that the tools used for genus assignment match their biological assumptions to avoid misguided inferences.

## Background

Bacterial taxonomy is the classification of bacterial strains into lineages ranging from phyla to species within the domain Bacteria. This is critical to our understanding of bacterial diversity by creating a coherent framework that reflect their evolutionary relationships. Two elements show that accurate taxonomic placement of microorganisms is more important now than ever: (i) a very high fraction of bacteria, both in the environment and host-associated microbiomes, remain to be described and named (Sutcliffe, Rosselló-Móra, and Trujillo 2021; Thomas and Segata 2019); (ii) large-scale metagenomic studies in the last decade and now high-throughput cultivation methods are accelerating the pace of bacterial discovery (Hugenholtz et al. 2021). It is therefore essential to consolidate the system for classifying bacteria. Whilst bacteria had been classified based on morphology and phenotypic parameters for decades, the advent of genomics has revolutionised the way we classify bacteria, giving rise to a plethora of Overall Genome Relatedness Indices (Chun and Rainey 2014). However, in the process of classifying bacteria, it is essential to provide a robust framework that includes easy-to-implement parameters to classify organisms at each level of bacterial lineage, including the genus level, which is the focus in this article.

The Genome Taxonomy Database (GTDB) has proposed a standardized approach to group publicly available genomes into species clusters (Donovan H. Parks et al. 2020), resulting in an invaluable resource that is regularly updated and adopted by the community (Donovan H. Parks et al. 2022). The approach is resilient to genome contamination, which can plague public repositories (Mussig et al. 2024), but also includes cases of taxonomic incongruence with previously described and accepted species names (Donovan H. Parks et al. 2020).

In the last three years alone, the number of bacterial genomes in the RefSeq collection has increased by 35,000 per year, both from isolates and metagenomes (Haft et al. 2024). To analyse these genomes, we need clear and rapid methods for taxonomic assignment. For species, Average Nucleotide Identity (ANI, Jain et al. 2018) has been developed and shown to delineate species almost unanimously (Donovan H. Parks et al. 2020). However, ANI is unable to delineate genera (Qin et al. 2014). While no single ANI threshold has been proposed for genus delineation, family-specific thresholds have been suggested using both the ANI value and alignment fraction, however the lack of a universal threshold limits their usability (Barco et al. 2020). An alternative to ANI is the Average Amino acid Identity (AAI) which uses protein sequences instead of genomic nucleic sequences (Konstantinidis and Tiedje 2005). A genus delineation threshold based on AAI has been proposed (Konstantinidis, Rosselló-Móra, and Amann 2017), but this approach is rarely applied (Dieckmann et al. 2021; Kim, Park, and Chun 2021; Medlar, Törönen, and Holm 2018). A protein sequence-based genus delineation method proposed with an interpretable metric is the Percentage of Conserved Proteins (POCP) (Qin et al. 2014). In this system, if two bacterial entities share more than half of their conserved proteins, i.e., POCP > 50%, they are considered to belong to the same genus.

POCP is widely used in the community to assign novel bacterial taxa to known genera, or support the proposal of novel genera (Afrizal et al. 2022; Chaplin et al. 2020; González et al. 2020; Thomas C. A. Hitch et al. 2024; Wylensek et al. 2020; Kuzmanović et al. 2022; Liu et al. 2021; Orata et al. 2018; Sereika et al. 2023) However, the validity of the threshold value aforementioned has not been widely tested. In addition, a major limitation of POCP is that comparing all proteins within each genome to each other is computationally demanding. Given that the number of valid genus names almost doubled since the original proposition of POCP by Qin et al. Qin et al. (2014) (Figure S1), scalable methods as well as a timely re-evaluation of the POCP approach is needed.

Hernández-Salmerón et al. (2020) compared proteins alignment tools to find faster alternatives to find reciprocal best hits, without a loss in precision. They found that DIAMOND (Buchfink, Reuter, and Drost 2021), when switched to sensitive parameters instead of defaults, correctly found 87% of the reciprocal best hits of BLASTP (Camacho et al. 2009) in less than 8% of the computing time. Recently, Hölzer (2024) suggested the use of DIAMOND with ultra-sensitive settings to compute POCP faster than with BLASTP. This method was based on previously available code (Hölzer 2020), implemented as a nextflow workflow (Hölzer 2024). However, this new method was only validated on 5 genera, with 15 to 167 genomes each. Fundamental changes, such as the tool selected, can have significant effects on calculations, especially if 13% of matching proteins may not be found. Current implementations of POCP have also modified the calculation of POCP by limiting the calculation of conserved proteins to unique matches (Hölzer 2020, 2024; Riesco and Trujillo 2024; Lin 2021), in contrast to the original implementation of POCP which had no such limitation (Qin et al. 2014). These studies clearly show the urgent need for a clear definition of POCP to avoid divergent assumptions in tools between microbiologists and developers. Furthermore, there is an urgent need for extensive benchmarking of POCP to enhance reproducibility and accuracy given the increased number of genomic resources available.

To achieve this, we created a scalable alternative to BLASTP for computing POCP. This alternative is fast and did not compromise the accuracy of the original approach. Secondly, because POCP provides an interpretable and used metric but the proposed threshold needs to be assessed in a comprehensive manner, we evaluated the ability of the optimised POCP implementation to correctly delineate bacterial genera based on 2,358,466 pairwise comparisons of 4,767 genomes across 35 families.

## Results

### Finding a BLASTP alternative for POCP

First, we set out to identify a scalable alternative to BLASTP to compute the Percentage Of Conserved Proteins (POCP) to delineate genera. We evaluated ten proteins alignment methods (Table 4) based on three tools – BLASTP (Camacho et al. 2009), DIAMOND (Buchfink, Reuter, and Drost 2021) and MMseqs2 (Steinegger and Söding 2017).

We used GTDB genomes from a wide range of bacterial phylogenetic diversity across four phyla: *Bacillota* (6 families, 23 genera, 561 species), *Pseudomonadota* (6 families, 16 genera, 333 species), *Bacteroidota* (3 families, 7 genera, 217 species) and *Actinomycetota* (2 families, 6 genera, 124 species). We processed these 1,235 genomes and conducted 1,412,040 pairwise comparisons with a total of 32,316 CPU-Hours (3.7 in years). All the methods tested were faster than the reference BLASTP (Table 1).

**Table 1:** Fold change of computing metrics for the ten methods used in the benchmark compared to the BLAST_BLASTP method. The metrics include processing time as real-time, memory usage, CPU usage and disk usage as input/output (I/O). A fold change below 1 means the metric was lower, above 1 means it was higher, compared to the reference. The fold change values are median computed over n = 141,204 number of processes tracked per approach.

| Method | Time | Memory | CPU | Disk usage (I/O) |
| --- | --- | --- | --- | --- |
| BLAST |  |  |  |  |
| BLAST_BLASTP | 1.000 | 1.000 | 1.000 | 1.000 |
| BLAST_BLASTPDB | 0.429 | 6.978 | 3.991 | 0.640 |
| DIAMOND |  |  |  |  |
| DIAMOND_FAST | 0.000 | 4.142 | 4.196 | 1.644 |
| DIAMOND_SENSITIVE | 0.062 | 10.095 | 4.498 | 1.902 |
| **DIAMOND_VERYSENSITIVE** | **0.043** | **9.844** | **5.178** | **2.290** |
| DIAMOND_ULTRASENSITIVE | 0.156 | 29.795 | 4.901 | 4.943 |
| MMSEQS2 |  |  |  |  |
| MMSEQS2_S1DOT0 | 0.067 | 7.600 | 4.520 | 0.310 |
| MMSEQS2_S2DOT5 | 0.071 | 7.747 | 4.623 | 0.405 |
| MMSEQS2_S6DOT0 | 0.140 | 16.270 | 5.292 | 0.837 |
| MMSEQS2_S7DOT5 | 0.314 | 16.281 | 5.675 | 1.582 |

BLASTPDB, the database method of BLASTP (Table 4) that enables paralleled computations, was only half the time of BLASTP on average. In contrast, all DIAMOND and MMSEQS2-based methods ran in 20x and 11x the speed of BLASTP, at the cost of using more memory, CPU, and disk usage (Table 1). Thus, as expected, more sensitive methods consumed more resources in general.

An important criterion for a BLASTP alternative is to evaluate whether the accuracy of POCP calculation is comparable and not compromised for increased speed.

### DIAMOND gives POCP as accurate as BLASTP

BLASTPDB produced the exact same POCP values as BLASTP (Figure S2). The other methods did not perform as good. All methods of DIAMOND had a coefficient of determination (*_R_*^2^) above 0.99, except for DIAMOND_FAST that deviated from the expected values (Figure 1). All DIAMOND methods, especially DIAMOND_FAST, tended to underestimate POCP values (all dots were below the reference dashed line), meaning that they might assign genomes to different genera when they are from the same genus (top panels). Deviation from the BLASTP reference was aggravated when using the MMSEQS2_S1DOT0 method, mainly through underestimation (bottom panels). The other MMSEQS2 methods performed better, but still less good than the DIAMOND methods. They also tended to overestimate more than underestimate. All in all, the DIAMOND methods, especially with increased sensitivity, generated POCP values nearly as accurate as BLASTP for a fraction of the time, but we restrained from using the MMSEQS2 methods due to being less accurate.

**Figure 1:**
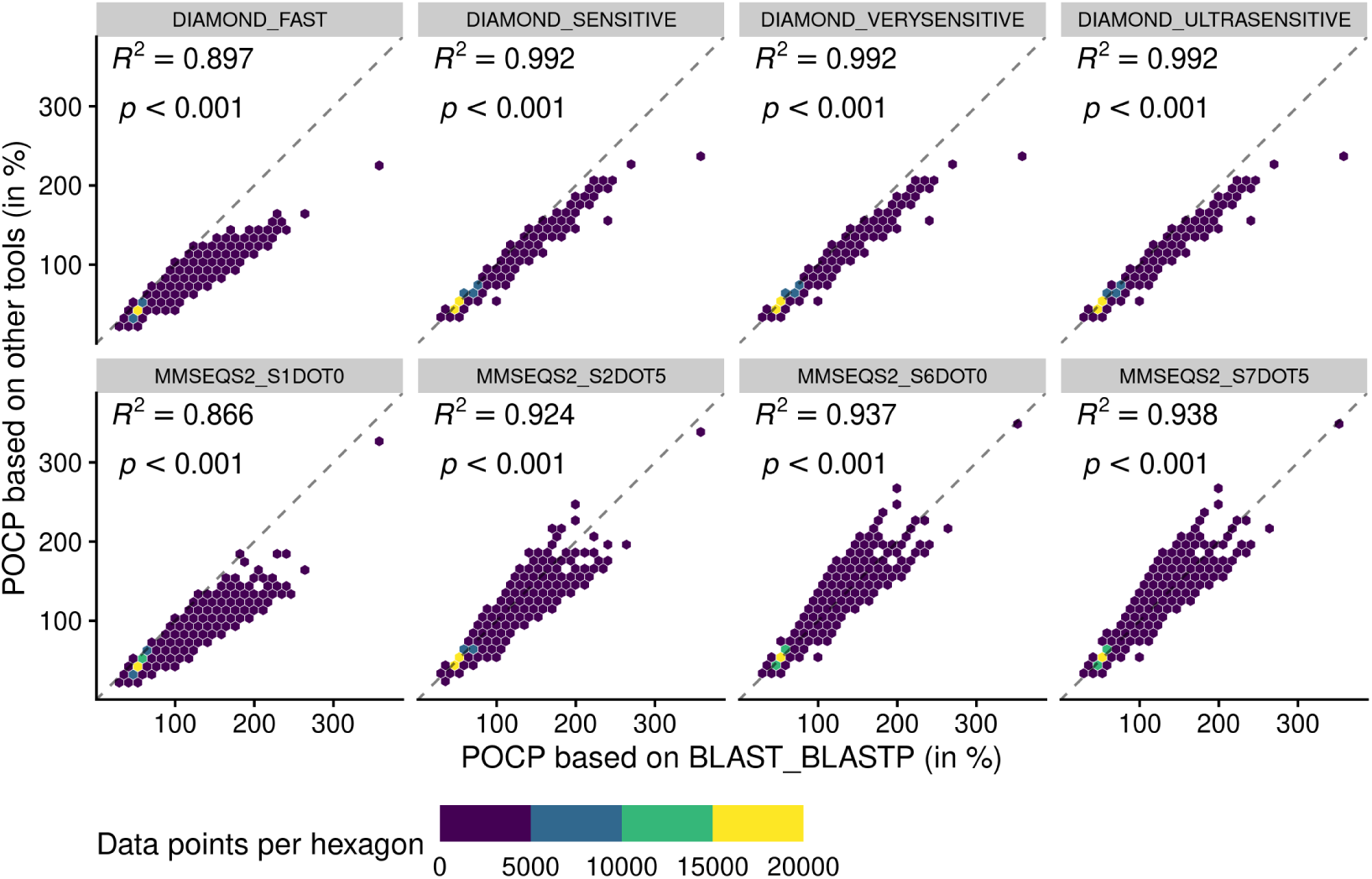
Adequacy between POCP values computed with the reference method BLAST_BLASTP and methods of faster alternatives: DIAMOND (Buchfink, Reuter, and Drost 2021) and MMSEQS2 (Steinegger and Söding 2017). Each point (n = 70,602 per tool) represents a POCP value between two genomes (see Equation 2). The colors represent the number of data points binned together in hexagons to avoid over-plotting. Coefficient of determination (*R*^2^) and associated p-value are shown on top of each linear regressions.

### Proposal for clear and fast computation of POCP values

We observed that all methods generated POCP values exceeding the supposed upper limit of 100%. Hence, we investigated the underlying reasons and suggest a clearer definition of POCP, termed POCPu.

During the alignment process, proteins sequences from the query can match multiple subject sequences in the case of duplicated genes. Whilst briefly mentioned in the original article that “The number of conserved proteins in each genome of strains being compared was slightly different because of the existence of duplicate genes (paralogs)” (Qin et al. 2014, 2211), the expected influence on the POCP values was lacking.

Therefore, we defined the POCP with unique matches (POCPu) between two genomes *Q* and *S* as:

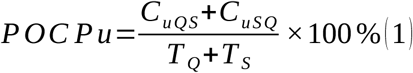

where *C_uQ S_* represents the conserved number of proteins from *the unique matches of Q* when aligned to *S* and, conversely, *C_u SQ_* the conserved number of proteins from *the unique matches of S* when aligned to *Q*; *T_Q_* +*T_S_* represents the total number of proteins in each of the two genomes being compared.

POCP values above 100% disappeared when using POCPu (Figure 2 and Figure S3). In general, the same patterns observed for POCP hold for POCPu, though with higher values of coefficient of determination (Figure 2 and Table 2). The 3 different sensitive methods of DIAMOND produced POCPu values that matched perfectly the ones produced by the reference method BLAST_BLASTP, with no underestimation as in the case of POCP previously. In contrast, the MMSEQS2 methods, whilst better with POCPu than POCP, still tended to underestimate POCPu values. All in all, POCPu is closer to BLASTP than POCP (Figure 2) thus guiding our choice to create an accurate BLASTP alternative.

**Figure 2:**
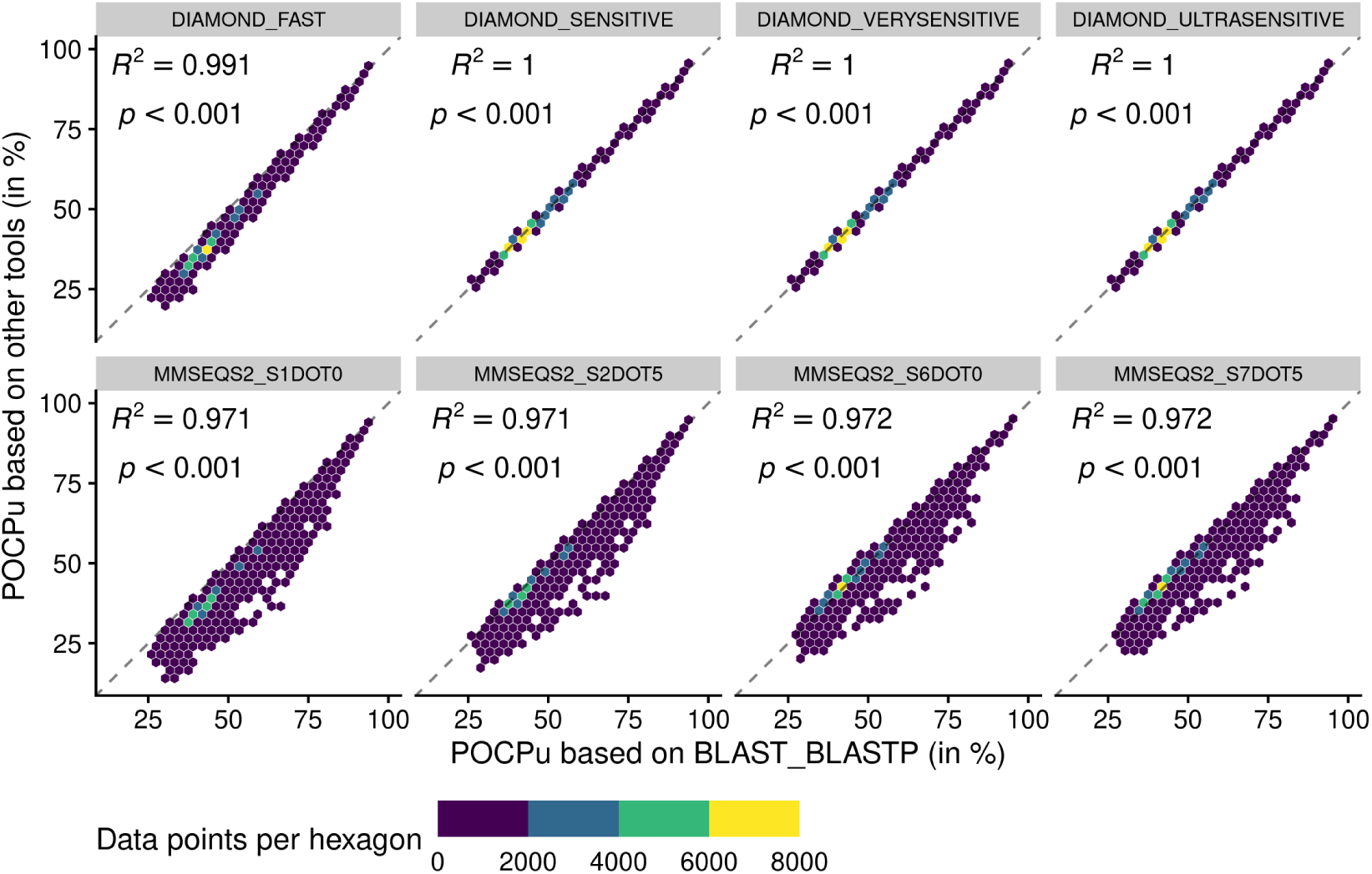
Adequacy between POCPu values computed with the reference method BLAST_BLASTP and methods of faster alternatives: DIAMOND (Buchfink, Reuter, and Drost 2021) and MMSEQS2 (Steinegger and Söding 2017). Each point (n = 70,602 per tool) represents a POCPu value between two genomes (see Equation 1). The colors represent the number of data points binned together in hexagons to avoid over-plotting. Coefficient of determination (*R*^2^) and associated p-value are shown on top of each linear regressions.

**Table 2:** Coefficient of determination (*R*^2^) and associated p-value for linear regressions matching the POCP and POCPu values computed by each method against the respective POCP and POCPu values of the reference method BLAST_BLASTP. Each linear regression are based on n = 70,602 comparisons per method. Methods are sorted by decreasing POCPu *R*^2^ values.

| Method | POCP |  | POCPu |  |
| --- | --- | --- | --- | --- |
| | $R^2$ | $p$-value | $R^2$ | $p$-value |
| BLAST_BLASTPDB | 1.00000000 | <0.001 | 1.00000000 | <0.001 |
| DIAMOND_ULTRASENSITIVE | 0.9920389 | <0.001 | 0.9997605 | <0.001 |
| DIAMOND_VERYSENSITIVE | 0.9919744 | <0.001 | 0.9997574 | <0.001 |
| DIAMOND_SENSITIVE | 0.9916398 | <0.001 | 0.9997438 | <0.001 |
| DIAMOND_FAST | 0.8967444 | <0.001 | 0.9913959 | <0.001 |
| MMSEQS2_S7DOT5 | 0.9379505 | <0.001 | 0.9722865 | <0.001 |
| MMSEQS2_S6DOT0 | 0.9369705 | <0.001 | 0.9720813 | <0.001 |
| MMSEQS2_S2DOT5 | 0.9242783 | <0.001 | 0.9713977 | <0.001 |
| MMSEQS2_S1DOT0 | 0.8659630 | <0.001 | 0.9713360 | <0.001 |

In summary, the BLAST_BLASTPDB method was a strong contender for a BLAST_BLASTP alternative because it performed exactly the same as BLAST_BLASTP (Figure S2, Figure S3 and Table 2) in twice the speed (Table 1), at the cost of using more resources. However, the DIAMOND sensitive methods were way faster with excellent adequacy with BLAST_BLASTP, especially for POCPu (Table 2). Whilst DIAMOND_ULTRASENSITIVE had the highest *R*^2^ value using POCPu (Table 2), it also had the highest memory consumption and disk usage (Table 1). A more sustainable alternative is DIAMOND_VERYSENSITIVE that performed 10 times faster than BLAST_BLASTPDB, in less than a twentieth of the time of the time of BLAST_BLASTP, while still maintaining reasonable usage of the computing resources (Table 1). More importantly, POCPu values calculated using DIAMOND_VERYSENSITIVE gave results that were extremely close to the reference BLAST_BLASTP (Figure 2 and Table 2) and were essentially identical to DIAMOND_ULTRASENSITIVE POCPu *R*^2^ up to 5 digits (Table 2). Therefore, we consider DIAMOND_VERYSENSITIVE to be a valid and scalable alternative to BLAST_BLASTP for POCP/POCPu computations.

### Unique matches enhance the accuracy of genus delineation

Next, we evaluated the 50%-threshold of POCP and POCPu to determine their reliability to delineate bacterial genera. This analysis was based on a broader range of genera to capture additional diversity: *Pseudomonadota* (15 families, 66 genera, 1,736 species), *Actinomycetota* (8 families, 27 genera, 1,584 species), *Bacteroidota* (6 families, 27 genera, 886 species) and *Bacillota* (6 families, 23 genera, 561 species). We included all the 4,767 genomes and calculated POCP and POCPu for 1,087,630 pairwise comparisons using the DIAMOND_VERYSENSITIVE method (Figure 3).

**Figure 3:**
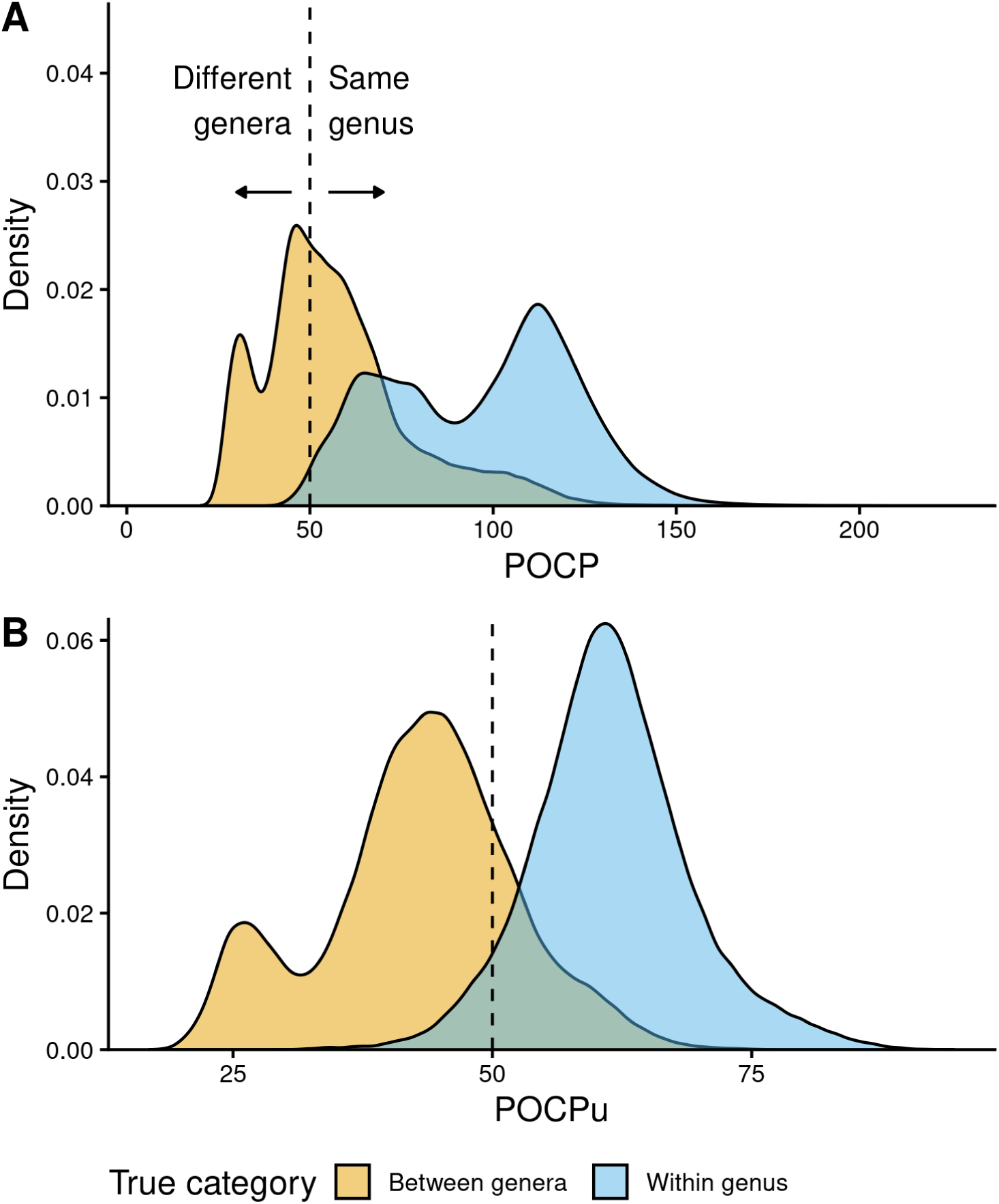
Distribution of POCP (A) and POCPu (B) values for all pairwise genome comparisons: Between genera (n = 321,189) in orange and Within genus (n = 222,626) in sky blue. The GTDB taxonomy was used as reference for the confusion matrix (true/false positives and negatives). POCP and POCPu values were calculated with our recommended method DIAMOND_VERYSENSITIVE (Table 4); they range from 20 to 236.9 for POCP and 16.9 to 94.6 for POCPu. The dashed lines indicate the standard 50% threshold for genus delineation.

Instead of two bell-shaped distributions, with the 50% threshold separating between-genera POCPs (left) from within-genus POCPs (right), we observed overlapping POCP distributions (Figure 3 A). This was associated with a high number of false positives (FP = 188,155), where between-genera values were >50%, especially compared with the number of true negatives (TN = 133,034), where between-genera values were <50%. Thankfully, most of the within-genus values were >50% (TP = 220,307), with only few below the threshold (FN = 2,319).

In contrast, POCPu was much closer to the expected results given the taxonomic assignments of each genome (Figure 3 B). Between-genera POCPu values followed a bi-modal distribution, with the highest peak and most of the distribution remaining below the 50%-threshold (TN = 253,860). Nonetheless, a fraction of between-genera values spilled over the threshold, representing false positives, i.e., different genera when they are not (FP = 188,155). As in the case of POCP, within-genus POCPu values were above the threshold of 50% (TP = 209,660), with a few below the threshold (FN = 12,966). All in all, considering only unique protein matches improves genus delineation.

To quantify these findings on the confusion matrix (true/false positives and negatives), we used the Matthews Correlation Coefficient (MCC, Equation 3), which is a binary classification rate that gives a high score only when the classifier correctly predicts most of positive and negative cases. POCPu (MCC = 0.72) surpassed POCP (MCC = 0.46) to delineate bacterial genera, which quantitatively confirmed the visual findings (Figure 3).

### Family-specific POCPu thresholds enable clearer genus delineation

Analysing the bacterial families separately questioned the universal threshold of 50% conserved proteins (Figure 4 and Figure S4). The large family of *Streptomycetaceae* (*Actinomycetota*), for instance, exhibited many false cases FN = 2,974 (2.8%) and FP = 7,110 (6.8%), despite even more true case TN = 1,231 (1.2%) and TP = 93,338 (89.2%), and consequently had a very low MCC (Figure 4 A). In 7 families out of 35, POCPu was clearly not adequate to delineate genus using a threshold of 50%, as indicated by low MCC (MCC *≤* 0.25; Figure 4 B). In contrast, POCPu delineated bacterial genera accurately for 18 families (MCC *≥* 0.7; Figure 4 B).

**Figure 4:**
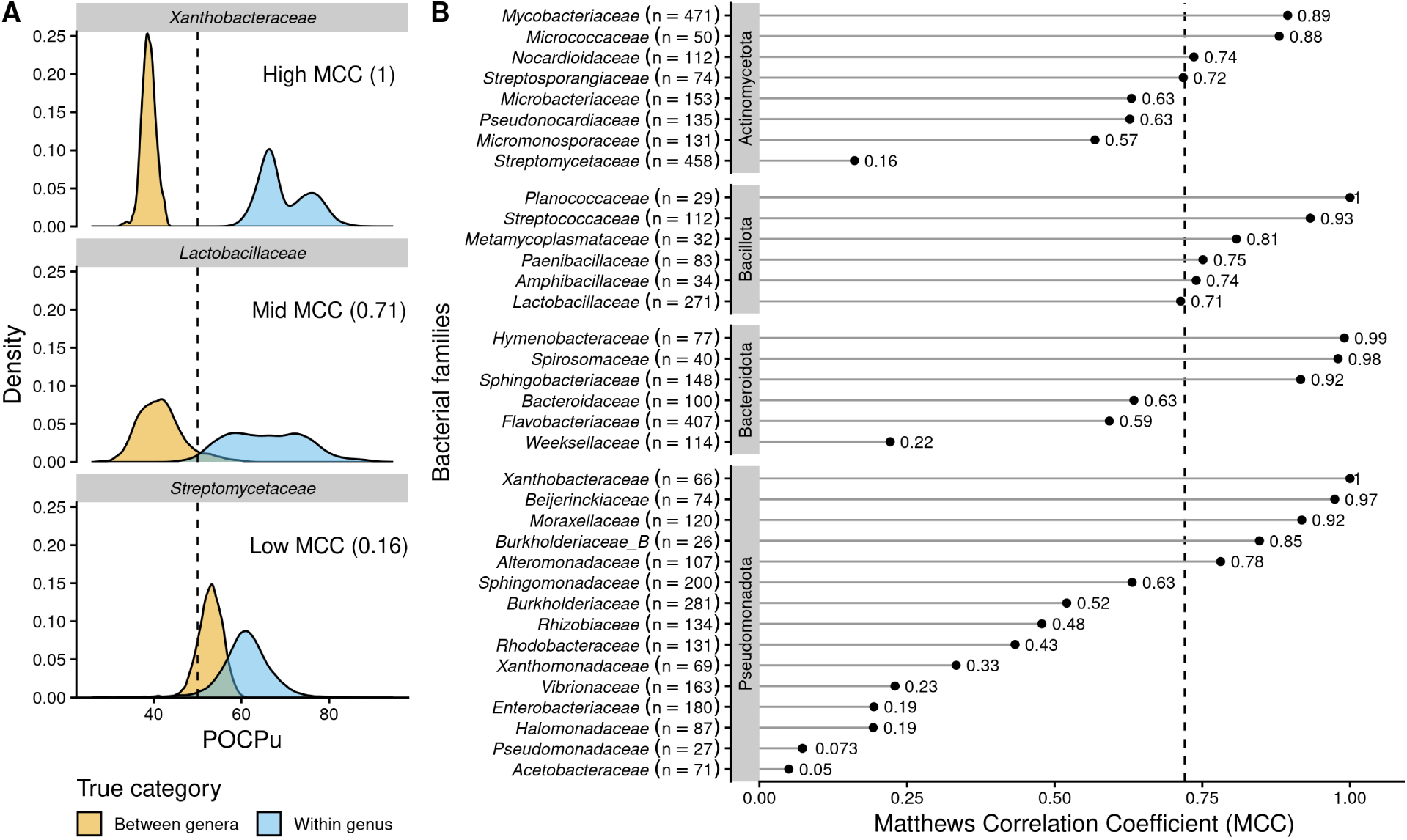
POCPu delineates bacterial genera in a family-specific manner. (A) Three representative examples of family-specific genus delineation capacity where POCPu values can (i) overlap and hamper genus delineation (top; example = Streptomycetaceae); (ii) be neatly distinct and allow for genus delineation (bottom; example = Xanthobacteraceae); or (iii) any scenario in between (middle; example = Lactobacillaceae). The dashed lines indicate the standard threshold of 50% conserved proteins. (B) The ability of POCPu to delineate genera was quantified for each of the 35 families analysed using the Matthews Correlation Coefficient (MCC, Chicco and Jurman 2020). An MCC of −1 and +1 indicates perfect misclassification or classification, respectively; random genus delineation corresponds to MCC = 0. The dashed line indicates the global MCC on the whole dataset, across all families. The number of genomes included per family are indicated in brackets next to the family names. The families were ranked per phyla (alphabetically; vertical facets in grey) and then by decreasing MCC within each phylum. The visual POCPu distributions for all families are provided in Figure S4.

Due to these differences between families, we looked for family-specific POCPu thresholds by maximizing MCC to improve genus delineation (Figure S4 and Table 3). With this procedure, thresholds other than the default 50% would enhance classification for 19 families out of the 35 families. The genus delineation of 8 families was improved, with at least 0.1-point increase in MCC (white squares in Table 3; Actinomycetota: *Streptomycetaceae* and *Streptosporangiaceae*; Bacillota: *Amphibacillaceae*, *Lactobacillaceae* and *Metamycoplasmataceae*; Pseudomonadota: *Acetobacteraceae*, *Burkholderiaceae_B* and *Vibrionaceae*). For 11 additional families maximum MCC above 0.7 were even obtained (black squares in Table 3; Actinomycetota: *Micromonosporaceae* and *Pseudonocardiaceae*; Bacteroidota: *Bacteroidaceae* and *Weeksellaceae*; Pseudomonadota: *Burkholderiaceae*, *Enterobacteriaceae*, *Halomonadaceae*, *Pseudomonadaceae*, *Rhizobiaceae*, *Rhodobacteraceae* and *Xanthomonadaceae*). Interestingly, in 2 cases, the optimal POCPu threshold was lower than the standard threshold: 43% for *Bacteroidaceae* (*Bacteroidota*) and 45.5% for *Metamycoplasmataceae* (*Bacillota*). In 17 cases, new thresholds higher than 50% conserved proteins better separated genomes from within genus and between genera (Table 3).

**Table 3:**
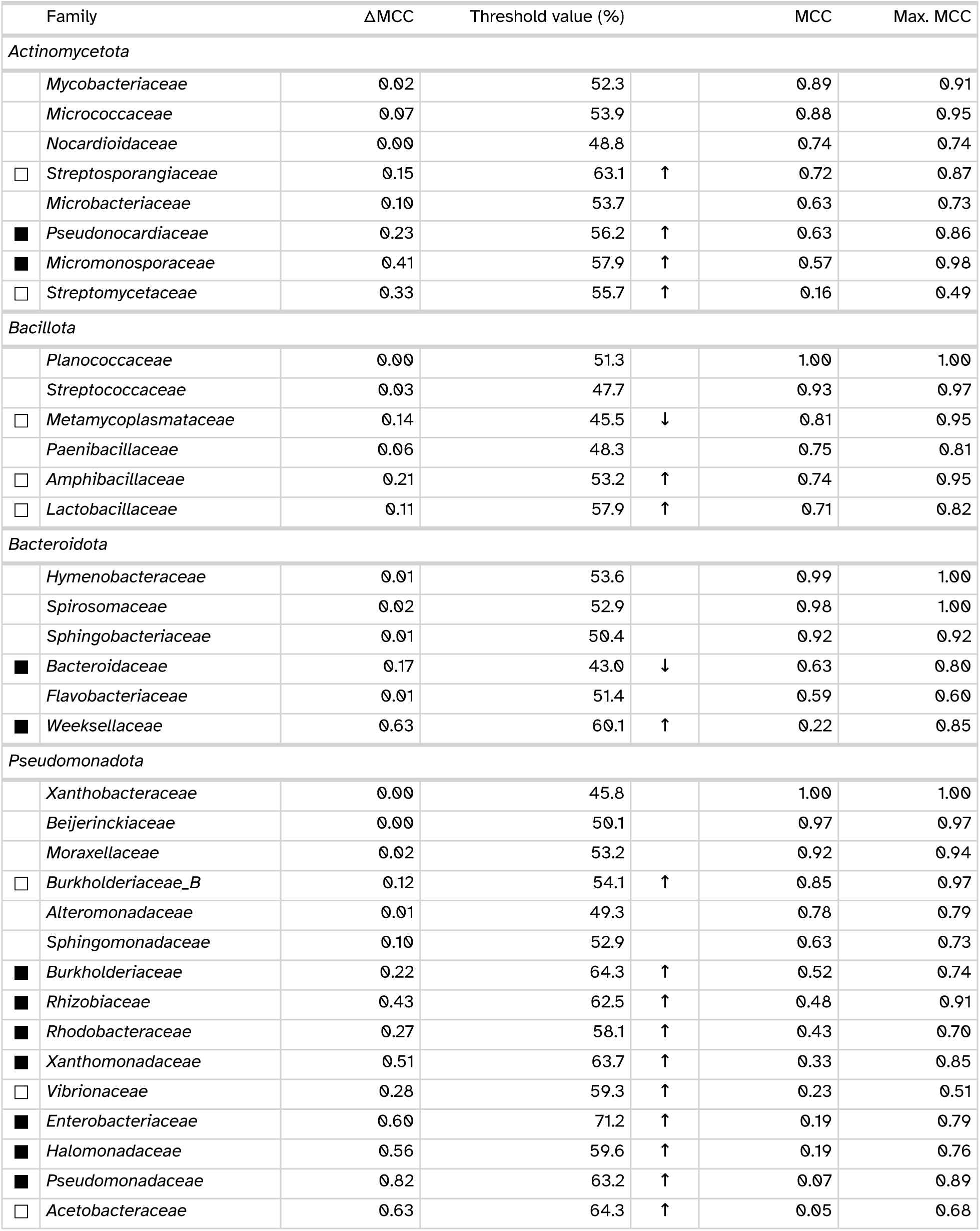
Proposal of family-specific POCPu thresholds for genus delineation. The thresholds were obtained after maximizing the MCC value to separate between-genera from within-genera distributions. Squares indicate that MCC value change was greater than 0.1 with the optimized threshold, filled squares denote rescued families from MCC < 0.7 to MCC > 0.7, whilst empty squares indicate improved genus delineation. Arrows highlight potential family-specific threshold worth considering to replace the default of 50% with the direction of change. Families without squares already delineate genera correctly with the default of 50%.

Because POCP was previously proposed to be influenced by genome size (Riesco and Trujillo 2024), we used the large pairwise comparisons dataset to assess whether the changes in threshold were linked to differences in genome size. If POCPu is influenced, we reasoned that its genus delineation power should also be influenced, therefore we expected stronger genome size differences in the families for which an alternative POCPu threshold was found. We found no evidence that POCPu is affected by differences in genome size (Figure S5 A) nor proteins number (Figure S5 B).

## Discussion

Qin et al. (2014) proposed to separate bacterial genera using the Percentage of Conserved Proteins (POCP) in genomes more than 10 years ago. The descriptions of many novel genera note POCP values oscillating around the proposed threshold value of 50%, suggesting it is not a clear-cut separation (Afrizal et al. 2022; Thomas C. A. Hitch et al. 2024; Wylensek et al. 2020). We therefore set out to re-evaluate genus-level delineation based on POCP using a comprehensive dataset, and to develop a faster and clearer method. We show that DIAMOND_VERYSENSITIVE can reliably replace BLASTP, speeding up the computing process by 20x. In addition, we addressed an assumption made in previous POCP implementations (e.g., Hölzer 2020, 2024; Lin 2021), and thereby clearly defined an alternative POCP metric – POCPu – that uses only unique matches, thereby making genus delineation more accurate.

Genus names occur before species names in the binomial nomenclature of bacteria, and are therefore an important first contact with bacterial entities. They are key to existing knowledge in databases or articles and provide intuitive information on the evolutionary history and ecological roles of the organisms under study (Reimer et al. 2022; Rosonovski et al. 2024; Schoch et al. 2020). The system works best when resources and tools follow FAIR principles (e.g., https://nfdi4microbiota.de, Wilkinson et al. 2016), as done in this work. We demonstrated that not all tools and parameters are suitable to speed up BLASTP; some combinations, whilst extremely fast, under-or overestimate POCP values, resulting in erroneous splitting or merging of genera. While our analysis included phylogenomically diverse taxa, it was limited to the taxa within GTDB (Donovan H. Parks et al. 2020; Donovan H. Parks et al. 2022) and focused on the dominant bacterial fraction. Despite the comprehensive nature of GTDB, our final genome selection might have missed important taxonomic groups of interest to readers. However, filtering the studied taxa was necessary to obtain enough data points per taxa to ensure statistical robustness. Previous studies on many-vs-many proteins alignment comparison used less phylogenetic diversity: 4 genomes from 4 genera in Hernández-Salmerón and Moreno-Hagelsieb (2020), maximum 167 genomes from 5 genera in Hölzer (2024). Riesco et al. Riesco and Trujillo (2024) evaluated much more genomes – 1,573 type strains – but calculated POCP – using Bio-Py (Lin 2021) – only to be compared with AAI and not to evaluate genus delineation.

Threshold-based approaches are always a matter of compromise, and do not provide one-size-fits-all solution. Regarding species delineation, Parks et al. (2022) stated: “The use of ANI to delineate species despite the lack of clear evidence for discrete species boundaries in the GTDB dataset is a pragmatic approach for organizing the rapidly growing biodiversity being discovered with metagenomic approaches” Donovan H. Parks et al. (2022). We share their vision and propose the use of POCPu as an interpretable and pragmatic approach to delineate bacterial genera. In an effort to improve this process, we suggest refining the classification by applying family-specific POCPu thresholds, as shown previously for *Rhizobiaceae* (Kuzmanović et al. 2022). However, one should only deviate from the standard threshold of 50% if the benefit is greater than the risk of creating more confusion. We have provided tentative thresholds for several families, for which confidence was high. Finally, it is important to remember that accurate taxonomic placement is best achieved when multiple lines of evidence are considered, as implemented in Protologger (Thomas C. A. Hitch et al. 2021). In the case of genera, POCPu decisions can be supported, for example by assessing the topology of phylogenetic trees, considering 16S rRNA gene identities (Yarza et al. 2014), and the result of GTDB-Tk analysis (Chaumeil et al. 2022).

### Conclusions

Our research contributes significantly to the field of genome-based microbiology by re-evaluating genus-level delineation based on the Percentage Of Conserved Proteins (POCP) proposed 10 years ago. We introduce a refined definition of POCP using only unique matches (POCPu) that improves genus delineation, and demonstrate the effectiveness of DIAMOND_VERYSENSITIVE as a faster yet accurate replacement for BLASTP. Robust methodologies like the one we put forward are needed as genomics has revolutionized bacterial classification. Finally, accurate bacterial taxonomy at scale is of critical importance in the era of rapid discovery of microorganisms.

## Methods

### Standardisation of protein sequences and taxonomy via GTDB

As GTDB provides curated taxonomy along with genomes and genome-derived proteins sequences (Donovan H. Parks et al. 2022), we used it as a reliable source of high-quality data in our benchmark. We used inclusion criteria to facilitate the selection of a diverse range of taxonomic groups from GTDB (r214) (N = 394,932 bacterial genomes), while maintaining achievable comparisons with the time, human, and computing resources available: (1) the bacteria had a valid name according to the List of Prokaryotic names with Standing in Nomenclature (Parte et al. 2020) and a representative genome was available (N = 11,699), (2) they belonged to a family with at least two genera (N = 5,904), and (3) to a genus with at least ten genomes (N = 4,767). Based on these criteria, the protein sequence files for the shortlisted bacteria were obtained from GTDB (Additional file 1).

### Definition of Percentage of Conserved Proteins (POCP)

The percentage of conserved proteins (POCP) between two genomes *Q* and *S* is defined as:

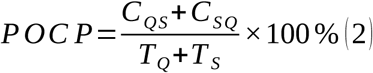

where *C_QS_* represents the conserved number of proteins from *Q* when aligned to *S* and conversely *C_SQ_* represents the conserved number of proteins from *S* when aligned to *Q*; *T_Q_*+*T_S_* represents the total number of proteins in each of the two genomes being compared (adapted from Qin et al. 2014). The range of POCP is theoretically [0; 100 %). Conserved proteins are defined as protein sequences matches from the query with an e-value < _10_*^−^* ^5^, a sequence identity > 40%, and an aligned region > 50% of the query protein sequence length.

Pairwise comparisons between a query sequence (*Q*) and a subject sequence (*S*) were defined by banning self-comparisons (*Q ≠S*) and considering reciprocal comparisons (*Q* - *S* and *S* - *Q*) only within the same family to avoid unnecessary expansion of the comparison landscape.

### Benchmarking methods for proteins sequence alignment

In Qin et al. (2014), guidance was provided on how to implement the computation of POCP, including the use of BLASTP. As ‘standard’ POCP method, we used BLASTP v2.14.0+ (Camacho et al. 2009) with parameters from Qin et al. (2014) (Table 4). We also considered a modified implementation of the BLASTP method, named BLASTPDB where BLAST databases are first built for the two genomes considered (Table 4). This allows parallel alignments on multiple CPU, which is not possible with BLASTP. We then included two tools that were designed as faster local-protein-alignment methods and used as alternatives to BLASTP: DIAMOND v2.1.6 (Buchfink, Reuter, and Drost 2021) and MMseqs2 v15.6f452 (Steinegger and Söding 2017). The former is used in Hölzer (2024) while the latter is used in EzAAI (Kim, Park, and Chun 2021). Similar to BLASTPDB, these methods require that a protein database is built for each genome before performing the alignment (Table 4). DIAMOND and MMseqs2 were both used with four different sensitivity thresholds proposed recently (Table 4) (Buchfink, Reuter, and Drost 2021).

**Table 4:** List of the ten methods and associated parameters for the many-versus-many proteins alignments tools used in the benchmark. The recommended approach is indicated in bold.

| Method | Database creation | Parameters |
| --- | --- | --- |
| BLAST |  |  |
| BLAST_BLASTP | No | --evalue 0.000001 --qcov_hsp_perc 50.0 |
| BLAST_BLASTPDB | Yes | --evalue 0.000001 --qcov_hsp_perc 50.0 |
| DIAMOND |  |  |
| DIAMOND_FAST | Yes | --evalue 0.000001 --query-cover 50.0 --fast |
| DIAMOND_SENSITIVE | Yes | --evalue 0.000001 --query-cover 50.0 --sensitive |
| **DIAMOND_VERYSENSITIVE** | **Yes** | **--evalue 0.000001 --query-cover 50.0 --very-sensitive** |
| DIAMOND_ULTRASENSITIVE | Yes | --evalue 0.000001 --query-cover 50.0 --ultra-sensitive |
| MMSEQS2 |  |  |
| MMSEQS2_S1DOT0 | Yes | --e-profile 0.000001 --cov-mode 1 -c 0.50 -s 1.0 |
| MMSEQS2_S2DOT5 | Yes | --e-profile 0.000001 --cov-mode 1 -c 0.50 -s 2.5 |
| MMSEQS2_S6DOT0 | Yes | --e-profile 0.000001 --cov-mode 1 -c 0.50 -s 6.0 |
| MMSEQS2_S7DOT5 | Yes | --e-profile 0.000001 --cov-mode 1 -c 0.50 -s 7.5 |

All proteins matches were filtered to only keep matches with > 40 % identity to all the query sequences matches for POCP (*C_QS_* and *C_SQ_* in Equation 2) and only unique query sequence matches for POCPu (*C_QS_* and *C_SQ_* in Equation 1). The filtering was adapted to the method as the range of percentage of identity in MMseqs2 is [0 *−*1) and [0 *−*100) for BLAST and DIAMOND. The total number of proteins per genomes (*T_Q_* and *T _S_* in Equation 2 and Equation 1) was computed using seqkit stats v2.2.0 (Shen et al. 2016).

Linear regressions were implemented using R version 4.3.1 (2023-06-16) to fit the expected POCP (or POCPu) values obtained via the BLAST_BLASTP reference method against the other methods considered. The coefficient of determination R^2^ of the linear regression is used as an interpretable and bounded goodness-of-fit measure between the expected and measured values instead of other errors measurement (Chicco, Warrens, and Jurman 2021). We did not rely on the adjusted coefficient of determination as the linear regressions had only one predictor, namely the POCP (or POCPu) values of the evaluated method.

### Metrics to evaluate genus delineation

We computed classification metrics with a positive event defined as “both genomes belong to the same genus”. Thus, for a pair of bacterial genomes with a POCP (or POCPu) >50%, the pair was a True Positive (*T P*) if it belonged to the same genus, else it was considered as a False Positive (*F P*). Conversely, for a pair of bacterial genomes with a POCP (or POCPu) *≤* 50%, the pair was a False Negative (*F N*) if it belonged to the same genus, else it was considered as a True Negative (*T N*). We then assessed the classification performance of both POCP and POCPu using Matthews Correlation Coefficient (MCC; Equation 3).

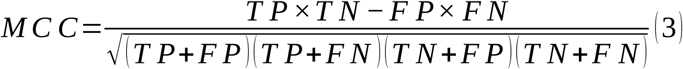

The coefficient has a range of [*−*1; +1) and is high in case of perfect classification, whilst a MCC of 0 indicates random classification. In addition, the MCC compensates for unequal class sizes compared to others metrics such as accuracy or F1-score (Chicco and Jurman 2020).

We used one dimensional optimization (Brent 1972) to find family-specific POCPu thresholds maximizing MCC values and separating between-genera from within-genera distributions. The optimization was run using the `optimize()` function from the R `stats` package (R Core Team 2023).

### Workflow implementation

Automatic protein sequences download, data pre-processing, many-versus-many protein alignments, POCP computation and delineation metrics calculations have been included in a workflow using nextflow v23.10.0 (Di Tommaso et al. 2017) workflow, based on components of nf-core (Ewels et al. 2020). The tools used are provided within Docker container (Merkel 2014) or bioconda (The Bioconda Team et al. 2018) environments to ensure reproducibility, scalability and ease future extensions of the present benchmarking work.

Nextflow natively keeps track of the time, CPU, memory and disk usage of each process in an execution trace log file, which we used to evaluate the computing resources utilization. Process duration is available as wall-time and real-time, the CPU usage is reported as a percentage of usage of a unique CPU, meaning multi-threaded processes will have a value higher than 100%.

Statistical analyses and visualization were conducted in R using targets v.1.7.0 (Landau 2021).

## Declarations

### Ethics approval and consent to participate

Not applicable

### Consent for publication

Not applicable

### Availability of data and materials

All the proteins sequences used in the analysis were downloaded from the GTDB (r214): https://data.gtdb.ecogenomic.org/releases/release214/214.0/genomic_files_reps/ gtdb_proteins_aa_reps_r214.tar.gz, along with the associated metadata: https://data.gtdb.ecogenomic.org/releases/release214/214.0/bac120_metadata_r214.tar. gz. The list of valid bacteria names was obtained from: https://github.com/thh32/Protologger/blob/0731adf80f1bbc5f8ee1904c8e9648ef45b13 303/DSMZ-latest.tab. Raw output files from the workflow are deposited at https://doi.org/10.5281/zenodo.14974869. Cleaned and formatted POCP/POCPu values and metadata tables for analysis are deposited at https://doi.org/10.5281/zenodo.14975029.

The Nextflow workflow used for the benchmark is available at https://github.com/ClavelLab/pocpbenchmark. Code for the analyses, figures and manuscript is available at https://github.com/ClavelLab/pocpbenchmark_manuscript.

### Competing interests

The authors declare that they have no competing interests.

## Funding

TC received funding from the German Research Foundation (DFG): project no. 460129525 (NFDI4Microbiota), and project no. 445552570.

## Authors’ contributions

CP and TCAH conceived the study with feedback from TC. CP performed the benchmark and analyzed the data with support from all authors. TC secured funding and provided computing resources. CP wrote the manuscript with inputs from all authors. All authors read and approved the final manuscript.

## Acknowledgements

The authors are grateful to Christian Schudoma, Daniel Podlesny and Mahdi Robbani for their help in fixing issues with an earlier version of the Nextflow workflow. CP thanks all members of the Clavel Lab for constructive feedback on the figures at the lab retreat 2024.

---

**Figure S1:**
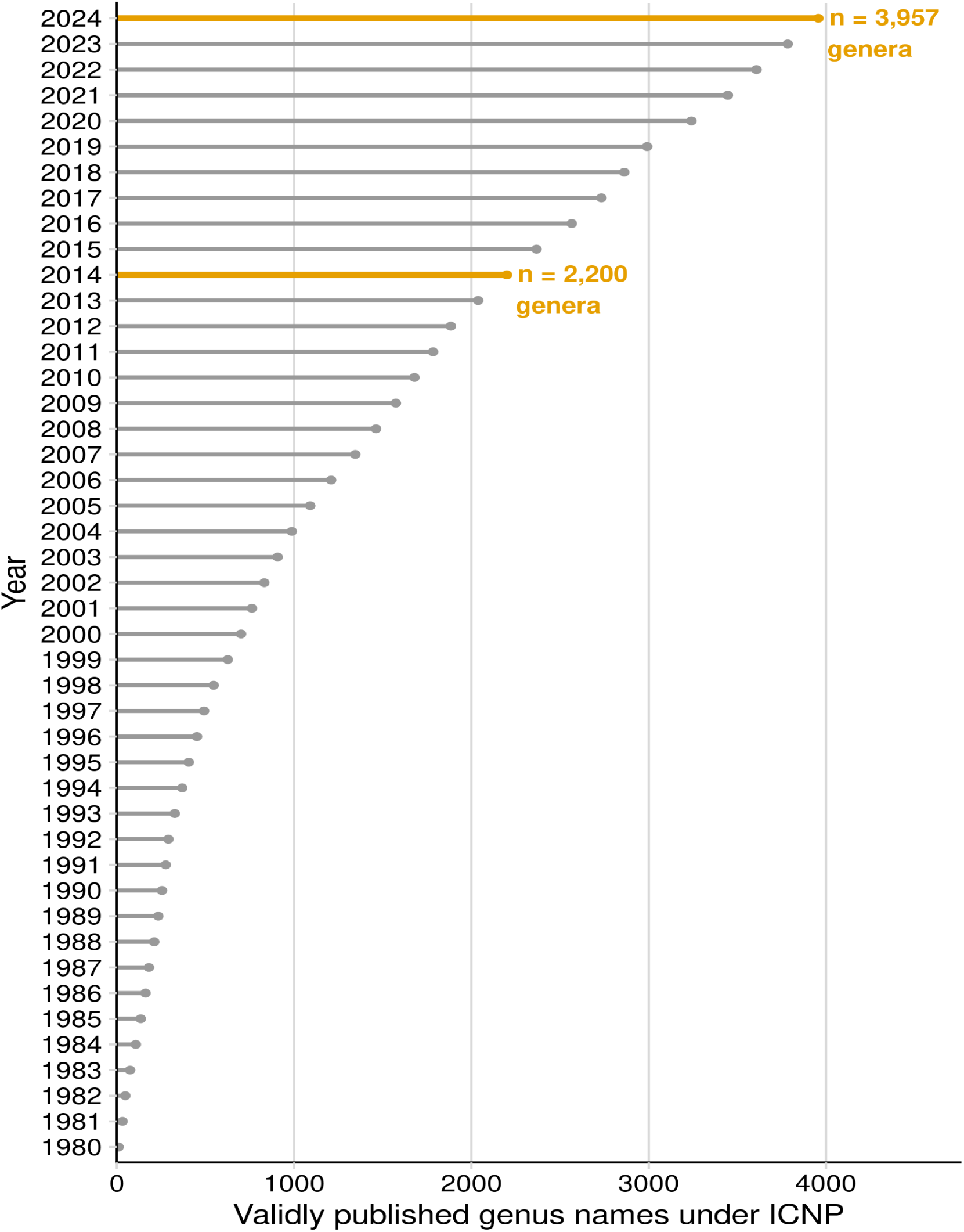
Cumulative number of validly published genera names according to the International Code of Nomenclature of Prokaryotes (ICNP). The year 2014 is highlighted as it corresponds to the year of publication of the paper by Qin et al. (doi: 10.1128/JB.01688-14) describing the Percentage of Conserved Proteins (POCP) to delineate genus. The number of valid genera is highlighted ten years later. The data was accessed on 2024-12-11 at the List of Prokaryotic names with Standing in Nomenclature (Parte et al. 2020).

**Figure S2:**
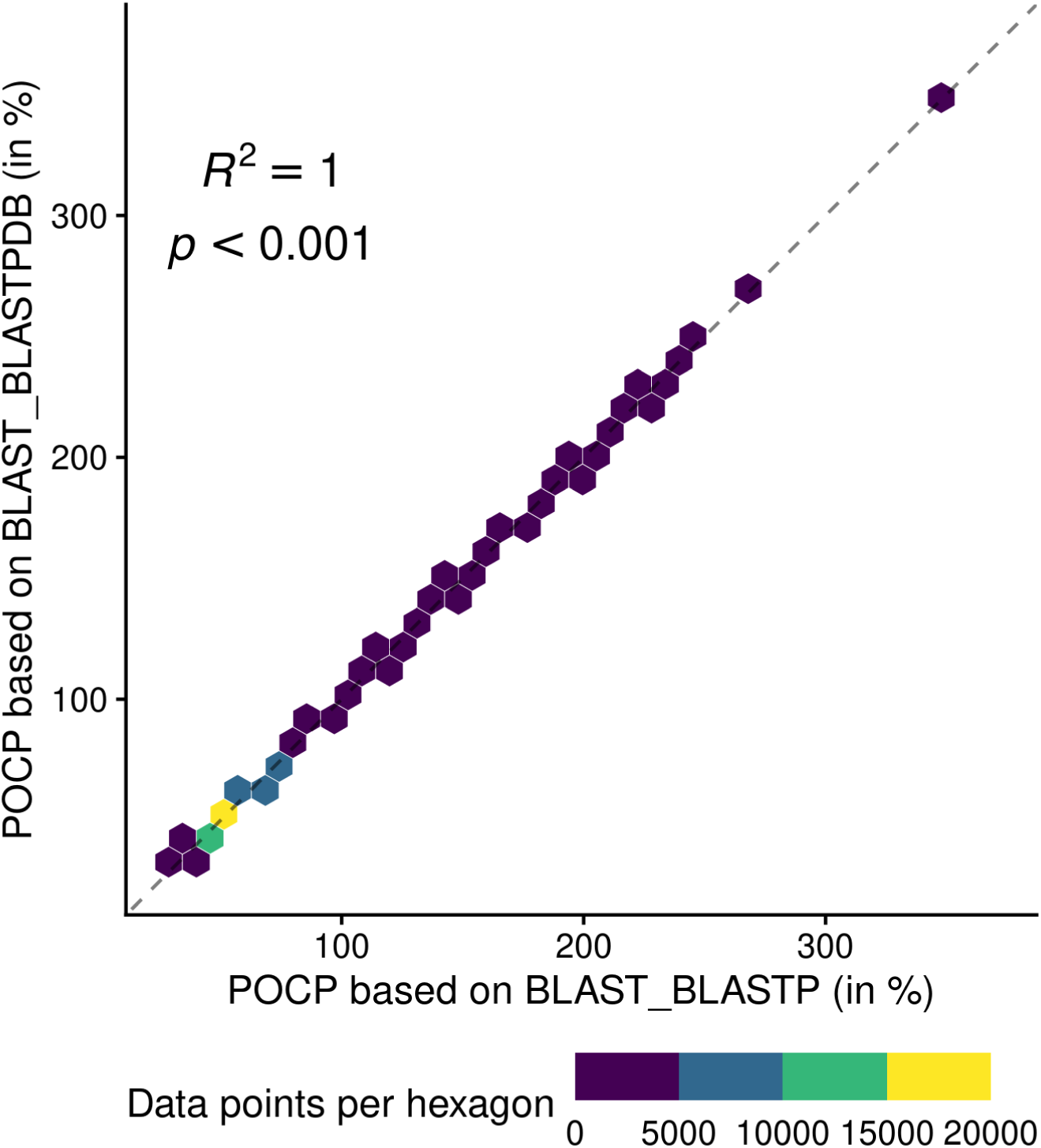
Adequacy between POCP values computed with the reference BLAST_BLASTP against the BLAST_BLASTPDB method that build databases before alignment. Each point (n = 70,602) represents a POCP value between two genomes (see Equation 2). The colors represent the number of data points binned together in hexagons to avoid over-plotting. Coefficient of determination (*R*^2^) and associated p-value are shown on top of each linear regressions.

**Figure S3:**
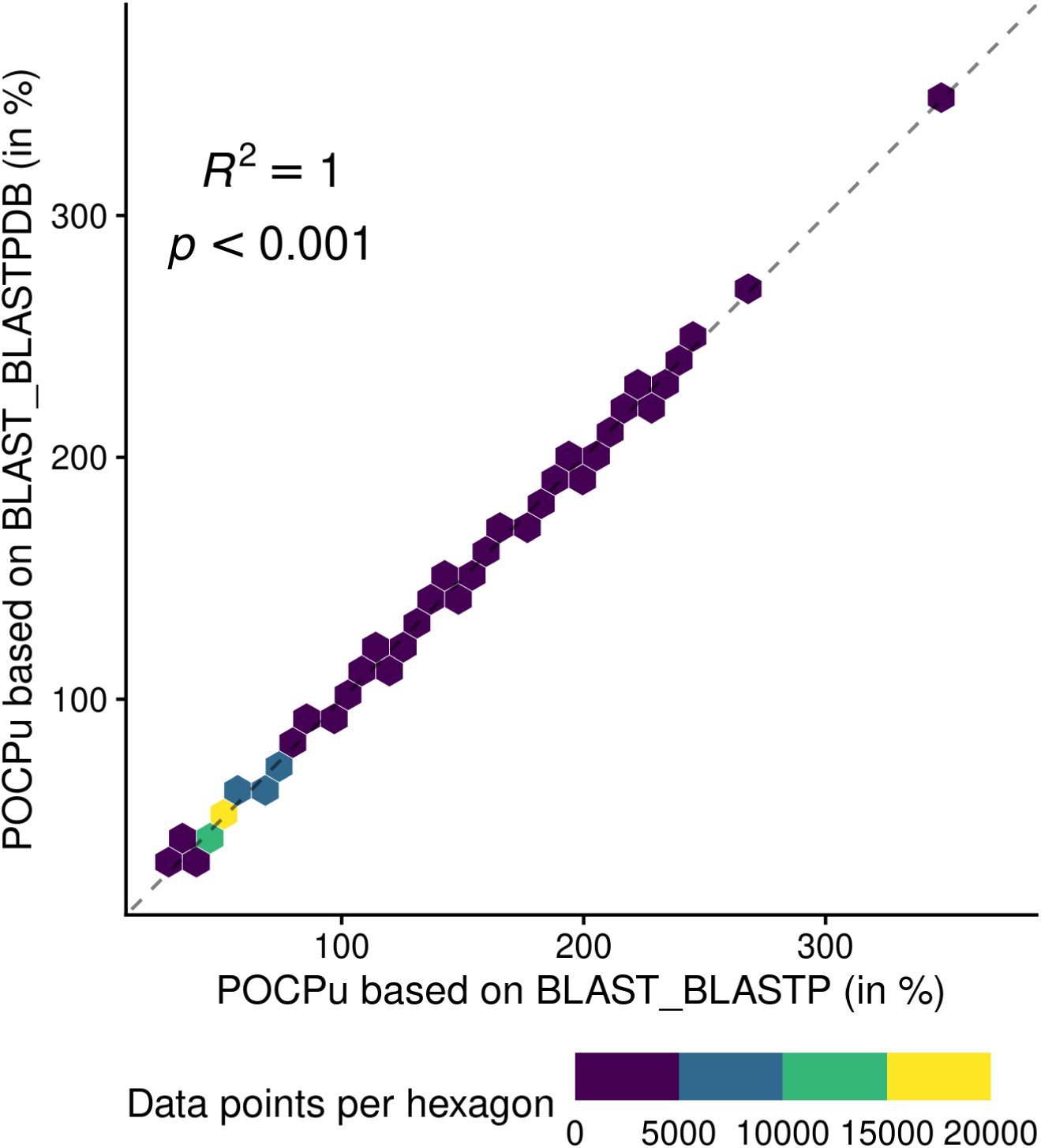
Adequacy between POCPu values computed with the reference BLAST_BLASTP against the BLAST_BLASTPDB approach that build databases before alignment. Each point (n = 70,602) represents a POCPu value between two genomes (see Equation 1). The colors represent the number of data points binned together in hexagons to avoid over-plotting. Coefficient of determination (*R*^2^) and associated p-value are shown on top of each linear regressions.

**Figure S4:**
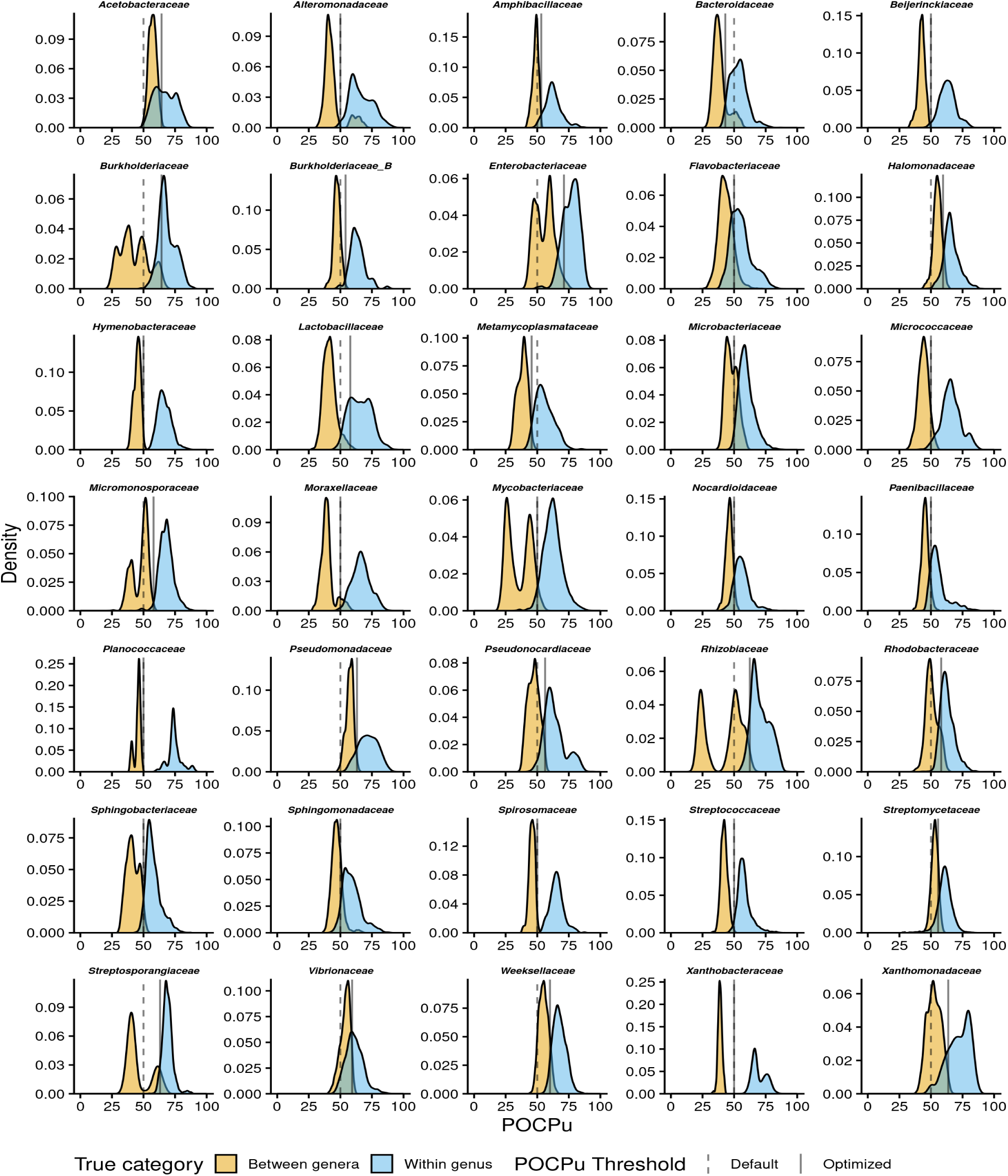
Distributions of POCPu values as in Figure 3 broken down per bacterial family. The true category is based on the GTDB taxonomy. The family-specific POCPu thresholds for genus delineation proposed in this study were taken from Table 3 and are indicated with plain vertical line, whilst the default POCPu threshold of 50% is indicated by dashed lines.

**Figure S5:**
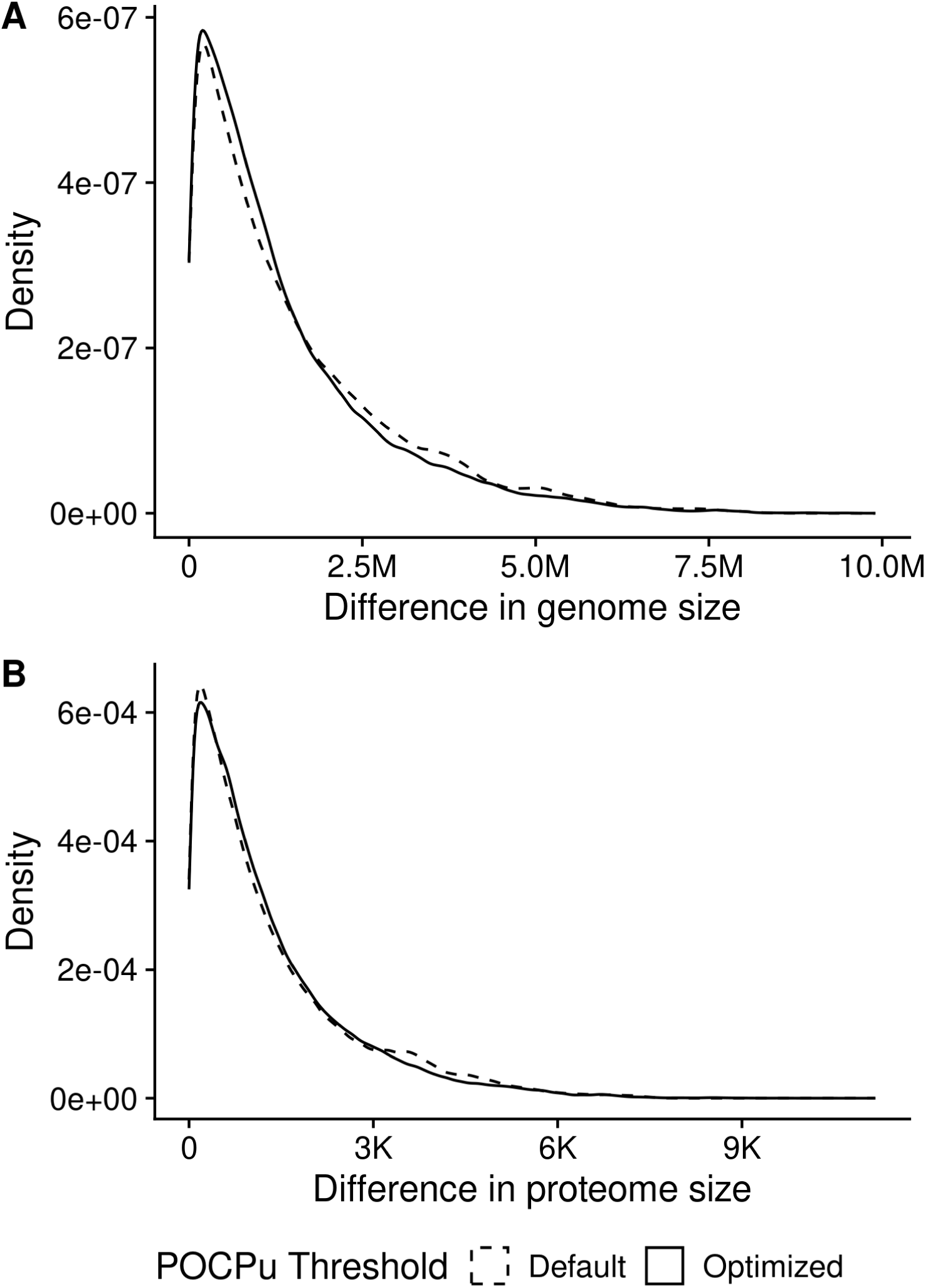
Distributions of differences in genome size (A) and proteome size (B) for families using default threshold or optimized thresholds. In case of an association between genome (or proteome) size and POCPu, we expected families for which optimized thresholds are proposed to have a shift towards larger differences explaining poor delineation performance in Figure 4 B. This was not the case, indicating that genome size and proteome size did not influence genus delineation. POCPu thresholds type were taken from Table 3.

## Additional files

- File name: Additional file 1
- File format: .csv
- Title of data: Shortlisted GTDB genomes for the benchmark
- Description of data: List of the accessions of Genome Taxonomy DataBase (r214), the type of benchmark in which they were included. The list includes the genome taxonomy from Domain to Species, summary statistics about the number of proteins, their cumulative, min, average and max length, as well as the genome size of the genomes. It is also available at https://github.com/ClavelLab/pocpbenchmark_manuscript/blob/920b847222e79511cdb6f08d41a7989cb5096961/Table_S1_shortlisted_genomes.csv

## Notes

https://doi.org/10.5281/zenodo.14974869

https://doi.org/10.5281/zenodo.14975029

https://github.com/ClavelLab/pocpbenchmark_manuscript/blob/920b847222e79511cdb6f08d41a7989cb5096961/Table_S1_shortlisted_genomes.csv

### Competing Interest Statement

The authors have declared no competing interest.

